# Selective Hydrolytic Defluorination of Branched Perfluorooctanoic Acid Isomers by a Haloacid Dehalogenase

**DOI:** 10.64898/2026.04.19.719434

**Authors:** Miao Hu, Shefali Bhardwaj, Sophia Newton, Alessandro T. Caputo, Michael J. Manefield, Colin Scott

**Affiliations:** CSIRO Environment, Black Mountain Science and Innovation Park, Canberra, ACT, Australia; CSIRO Advanced Engineering Biology Future Science Platform, Black Mountain Science and Innovation Park, Canberra, ACT, Australia; ARC Centre of Excellence in Synthetic Biology, ACT, Australia; UNSW Water Research Centre, School of Civil and Environmental Engineering, UNSW, Sydney, NSW, Australia; CSIRO Manufacturing, Research Way, Clayton, VIC, Australia

**Author notes:** Address correspondence to Miao Hu,; Colin Scott. Present email address (M.H.). School of Molecular Sciences, The University of Western Australia, Perth, WA 6009, Australia (C.S.).

**Keywords:** haloacid dehalogenase, hydrolytic defluorination, perfluorooctanoic acid (PFOA), branched isomers, enzyme promiscuity, PFAS

## Abstract

Per- and polyfluoroalkyl substances (PFAS) are highly resistant to enzymatic C–F bond cleavage, and hydrolytic defluorination of long-chain PFAS has rarely been demonstrated. Here, we report selective hydrolytic defluorination of branched perfluorooctanoic acid (PFOA) isomers by a haloacid dehalogenase (4A) from *Delftia acidovorans* strain D4B. A fluoride-specific riboswitch biosensor was used for initial substrate screening, followed by scaled-up assays in which fluoride release was quantified using a fluoride ion-selective electrode. Defluorination products were subsequently identified by liquid chromatography–mass spectrometry (LC-MS). Although purified 4A (10 μM) readily catalyzed hydrolytic defluorination of fluoroacetic acid, incubation of PFOA (0.5 mM) with purified 4A resulted in a statistically significant increase in fluoride release at elevated enzyme loading (500 μM). High-resolution LC-MS/MS analysis revealed that defluorination products originated from minor branched PFOA isomers rather than linear PFOA. Molecular docking analyses supported catalytically plausible binding geometries for branched PFOA isomers, positioning the substrate α-carbon within ∼4 Å of the catalytic aspartate residue. These findings demonstrate previously unrecognized hydrolytic reactivity of a haloacid dehalogenase toward branched PFAS isomers and expand the known catalytic scope of the haloacid dehalogenase family.

**GRAPHICAL ABSTRACT:** 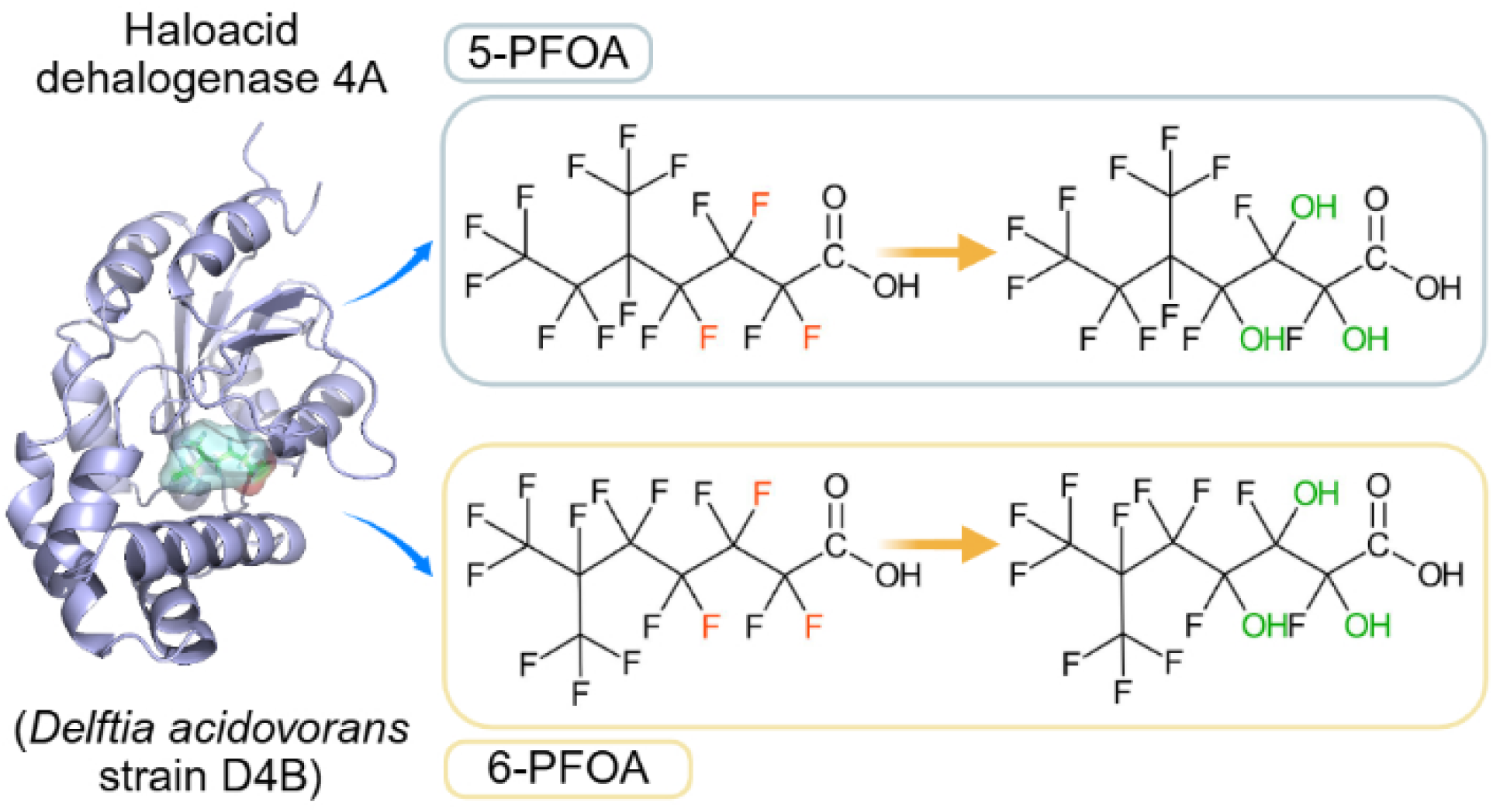

**SYNOPSIS:** Enzymatic defluorination of PFAS is rarely observed in environmental systems. This study identifies hydrolytic defluorination of branched PFOA isomers, improving understanding of PFAS defluorination at the enzyme level.

## INTRODUCTION

Per- and polyfluoroalkyl substances (PFAS) are synthetic organofluorine compounds characterized by exceptional chemical and thermal stability, resulting in their persistence and bioaccumulation across diverse environmental matrices.^1–3^ The extreme strength of the C–F bond is often used to explain why these compounds are resistant to natural degradation and accumulate in ecosystems.^4,5^ Although numerous bacterial and fungal strains capable of transforming or partially degrading PFAS have been reported.^6–9^ The enzymes responsible remain poorly understood. Only a small number of enzymes have been experimentally demonstrated to transform fluorinated carboxylic acids, either catalyzing C–F bond cleavage^10–12^ or the formation of CoA thioesters with aryl-CoA synthetases,^13–15^ highlighting plausible biochemical routes for PFAS transformation while leaving enzymatic mechanisms largely unresolved.

Among the enzyme families proposed to catalyze C–F bond cleavage, haloacid dehalogenases (HADs) have received sustained attention. HADs are hydrolytic enzymes that catalyze the substitution of halide ions in α-halogenated carboxylic acids via an S_N_2 displacement mechanism using a conserved catalytic aspartate (Asp) residue. Within this family, a subset of enzymes, commonly referred to as fluoroacetate dehalogenases, catalyzes hydrolytic C–F bond cleavage in fluoroacetate and related compounds, releasing fluoride and hydroxylated products. The biochemical mechanisms, structural features, and substrate specificities of HADs and fluoroacetate dehalogenases have been comprehensively reviewed elsewhere.^9^ Structural and mechanistic analyses indicate that HADs share a conserved catalytic architecture, including a nucleophilic Asp-centered catalytic motif and a flexible substrate-binding pocket. These features support activity toward a narrow range of α-halogenated carboxylic acids. However, despite this mechanistic versatility, direct biochemical evidence demonstrating HAD-mediated defluorination of environmentally relevant PFAS remains limited, particularly for long-chain or perfluorinated substrates.

In recent years, several studies have reported genomic and transcriptomic associations between HADs and PFAS transformation. Culture-based studies have identified bacterial strains capable of transforming long-chain perfluorocarboxylic acids. Genomic analyses of these strains revealed genes annotated as HADs, although no biochemical evidence has been provided to show a causal link.^16^ Similarly, metagenomic and metatranscriptomic analyses of PFOS-transforming consortia revealed elevated expression of genes annotated as HADs.^17^ Together, these observations indicate repeated associations between HADs and PFAS-impacted microbial systems. However, direct biochemical evidence for HAD-mediated C–F bond cleavage in environmentally persistent PFAS remains lacking.

Recently, a HAD (PZP66635.1) from *D*. *acidovorans* was investigated for PFOA defluorinase activity by expressing it in *Escherichia coli*. Cultures expressing P66635.1 and exposed to PFOA showed a modest, statistically insignificant, increase in fluoride release.^18^ Although this observation did not establish enzyme-mediated defluorination, it suggested that *Delftia*-derived HADs might merit further investigation. We therefore attempted to express and purify PZP66635.1. While we were unsuccessful, we achieved this using a homolog of PZP66635.1, HAD 4A from *Delftia acidovorans* strain D4B (25.26% sequence identity), which was isolated from a perfluorooctanoic acid (PFOA)-contaminated environment.^19^

In this study, we combined a fluoride-responsive riboswitch assay with biochemical analyses and structural modeling to evaluate the substrate scope of HAD 4A toward fluorinated compounds, including PFOA. We found that HAD 4A was active against the toxic natural fluorinated compound, fluoroacetate. Surprisingly, we also found that fluoride was released when HAD 4A was incubated with a PFOA solution. Careful analysis revealed that HAD 4A defluorinated two minor geometric isomers of PFOA (5-PFOA and 6-PFOA), rather than linear PFOA (the major component of the mixture). As far as we are aware, this is the first perfluorohydrolase activity observed in an enzyme.

## MATERIALS AND METHODS

### Materials

Fluoroacetic acid (95% purity) was obtained from Sigma-Aldrich (NSW, Australia). PFOA (98% purity) was obtained from SynQuest Laboratories (Florida, USA). Potassium fluoride (KF, Sigma-Aldrich) was used as a fluoride standard. *Escherichia coli* BL21 (DE3) was obtained from New England Biolabs. 5-Bromo-4-chloro-3-indolyl-β-D-galactopyranoside (X-gal) solution (20 mg/mL; Thermo Fisher Scientific, RO941) and 10 mM Tris buffer (pH 8.0, prepared from a 1 M stock; Thermo Fisher Scientific, AM9855G) were used for enzyme assays. Glycolic acid solution (70 wt% in H_2_O; Sigma-Aldrich, 420581) served as a non-fluorinated control substrate. All other reagents were of analytical grade and used as received.

### Protein expression and purification

The *E. coli* codon-optimized defluorinase gene encoding 4A was synthesized and cloned into the pET-28a(+) plasmid (GenScript, Singapore). This plasmid was used to transform *E*. *coli* BL21 (DE3) (New England Biolabs, Australia) via heat shock. Transformed cells were grown in LB medium containing kanamycin (50 μg/mL) at 37℃ until the optical density at 600 nm (OD_600_) reached 0.6–0.8. Protein expression was induced with 1 mM isopropyl β-D-1-thiogalactopyranoside (IPTG), and cultures were incubated for 24 h at 15, 20, 28, 30, and 37℃ with shaking at 200 rpm to assess heterologous protein production (Figure S1a). Proteins were separated by SDS-PAGE using NuPAGE^TM^ Bis-Tris 4–12% gels (Thermo Scientific) and 1×NuPAGE^TM^ MES SDS running buffer (Thermo Scientific) and stained with AcquaStain Protein Gel Stain (Bulldog Bio, NH, USA). SeeBlue Plus2 Pre-stained Protein Standard (Thermo Scientific) were used as molecular weight marker.

Large-scale expression and purification of 4A followed the procedures described previously^20^ except that an induction temperature of 20C was used. Purification used a Ni–NTA affinity chromatography and Superdex 200 size-exclusion chromatography on an ӒKTA fast protein liquid chromatography system (GE Healthcare). The purified HAD 4A was concentrated using a 10K molecular weight cutoff concentrator (Figure S1b). A portion was stored at −80°C for enzyme assays, with the remainder used for crystallization. The gene and protein sequences of 4A are provided in Table S1.

### Crystal structure

Purified HAD 4A (8 mg/mL) protein was screened against an array of sparse-matrix crystallization screens in sitting-drop vapor-diffusion trays (SwissSCI) set up on a Crystal Phoenix nanodispenser (ARI). Crystallization drops were set up with 150 nl protein and 150 nl crystallant and incubated at 293 K. Intersecting plate-like crystals of 4A appeared overnight in the Shotgun 2 screen^21^ in 2 M ammonium sulfate, 2% (v/v) polyethylene glycol 400, 0.1 M HEPES pH 7.5. 4A was cryprotected by adding 15% (v/v) ethylene glycol or glycerol to the reservoir for A4. The crystals were flash-cooled in liquid nitrogen. X-ray diffraction occurred at the Australian Synchrotron MX2 beamline tuned to 13 keV under a nitrogen vapor stream at 100 K.^22^ Diffraction images were processed with autoPROC utilizing XDS and the CCP4 suite programs Pointless, Aimless, Truncate.^23–26^ Phases were estimated with Phaser using AlphaFold2 generated models.^27,28^ Model building and refinement was carried out in Coot and autoBUSTER for 4A.^29–31^ A4 has 99.04% favored and 0% outliers in the Ramachandran plots. The crystal structure of 4A has been deposited in the Protein Data Bank (PDB) under the accession code 9C9E. Data collection, processing, and model statistics are summarized in Table S2.

### Docking

Ligand structures (fluorinated compounds) were obtained from PubChem when available. For compounds lacking three-dimensional (3D) coordinates, two-dimensional (2D) structures were generated in ChemDraw (version 23.1.1; Revvity Signals Software, Inc., Waltham, MA, USA), converted to three-dimensional conformations, and energy-minimized prior to docking.

The receptor (4A) and ligand structures were prepared using AutoDock Tools, and molecular docking simulations were performed with AutoDock Vina (v1.2.3).^32^ Docking poses were visualized in 3D using PyMOL (v3.1.4),^33^ and 2D protein–ligand interaction maps were generated using LigPlot+ (v2.3.1).^34^

### Defluorination assays

Scaled-up defluorination assays were conducted to evaluate enzymatic fluoride release from fluoroacetic acid and PFOA under defined conditions. All reactions were performed in a total volume of 5 mL using 10 mM Tris-HCl buffer (pH 8.0), with incubation conditions adjusted according to the experimental objective, as described below.

For fluoroacetic acid defluorination assays (Results Section 3.1), reactions contained purified 4A (10 μM) and fluoroacetic acid (1 mM). These reactions were incubated under the same conditions used for the plate-based biosensor assays (30℃, 200 rpm, 24 h). Control reactions contained an enzyme-only and a substrate-only to assess background fluoride levels.

For PFOA defluorination assays, experimental reactions consisted of purified 4A with PFOA at two enzyme concentrations: 10 μM 4A + 0.5 mM PFOA and 0.5 mM 4A + 0.5 mM PFOA. Negative controls included reactions containing purified 4A only (0.5 mM), PFOA only (0.5 mM), or buffer only. These reactions were incubated statically at room temperature (∼20℃) in the dark for up to 120 h. To verify enzyme activity under the high-enzyme conditions used for PFOA assays, positive control reactions were performed using purified 4A (0.5 mM) and fluoroacetic acid (0.5 mM) under identical incubation conditions. All defluorination assays were performed with two independent biological replicates, unless otherwise stated.

After incubation, a 100 μL aliquot from each reaction was used for plate-based riboswitch biosensor assays. The biosensor assay was performed following the protocol described in our previous work.^20^ The remaining reaction mixtures were subjected to fluoride quantification and LC/MS analysis to identify defluorination products.

### Analysis of defluorination products

#### Fluoride concentration measurement

Fluoride release was measured using a PerfectION^TM^ combined fluoride ion-selective electrode (Mettler Toledo, product no. 51344715) connected to a Seven2Go^TM^ S8 Pro ion meter (Mettler Toledo, Australia). Measurements were conducted in accordance with the manufacturer’s operating protocol, as previously described.^20^ Fluoride concentrations were determined at room temperature against a standard calibration curve prepared from freshly diluted KF solutions. Each assay included two biological replicates, and each sample was measured eight times as technical replicates to ensure analytical reproducibility.

#### Defluorinated product identification

Defluorination products from fluoroacetate and perfluorooctanoic acid (PFOA) were analyzed using a Q-Exactive HF mass spectrometer (Thermo Fisher Scientific) coupled with a U3000 UHPLC system, as described previously.^20^ Briefly, the instrument was operated in the negative electrospray ionization mode, and both full-scan and MS/MS modes were employed to identify and characterize the defluorinated intermediates. Chromatographic and MS parameters (solvent composition, gradient, and resolution) were identical to those previously reported. Spectral data were analyzed using Thermo Scientific XCalibur Qual Browser (version 4.1.50), and product identification was confirmed by matching theoretical and observed exact masses within ±10 ppm.

### Statistical analysis

Fluoride concentrations are reported as mean ± standard deviation (SD) of two biological replicates. Each biological replicate was measured in eight technical replicates. For each biological replicate, the mean value of the technical replicates was calculated before statistical analysis. Statistical differences in fluoride release from PFOA, with and without purified 4A, were evaluated using two-tailed unpaired Welch’s *t*-tests in GraphPad Prism v9.5.1 (GraphPad Software Inc.). A *P* value < 0.05 was considered statistically significant.

## RESULTS

### HAD 4A is a fluoroacetate defluorinase

We used a fluoride-specific riboswitch biosensor assay, developed previously,^20^ to test HAD 4A’s ability to defluorinate fluoroacetate. Incubation of purified 4A (10 µM) with fluoroacetic acid (1 mM) produced a clear blue signal in the plate-based assay, whereas no color development was observed in substrate-only controls (data not shown). This qualitative response indicated the release of fluoride from fluoroacetic acid and prompted further quantitative validation.

The HAD 4A-mediated fluoride release was assessed using a fluoride-selective electrode. Only background fluoride levels (1–3 µM) were detected in enzyme-only and substrate-only controls. In contrast, reactions containing both purified 4A (10 µM) and fluoroacetic acid (1 mM) yielded a markedly elevated fluoride concentration after 24 h, demonstrating enzyme-dependent C–F bond cleavage (Figure 1b). The expected defluorinated product was identified by mass spectrometry. Analysis of the reaction mixture revealed a dominant ion corresponding to glycolate, the expected hydrolytic product of fluoroacetic acid defluorination, with the observed *m/z* matching the theoretical value within <5 ppm error (Figure 1a). These results confirm that the detected fluoride originated from enzymatic cleavage of the C–F bond rather than nonspecific degradation. Full LC–MS/MS chromatograms, retention times, and fragmentation spectra are provided in Figure S2.

**Figure 1.**
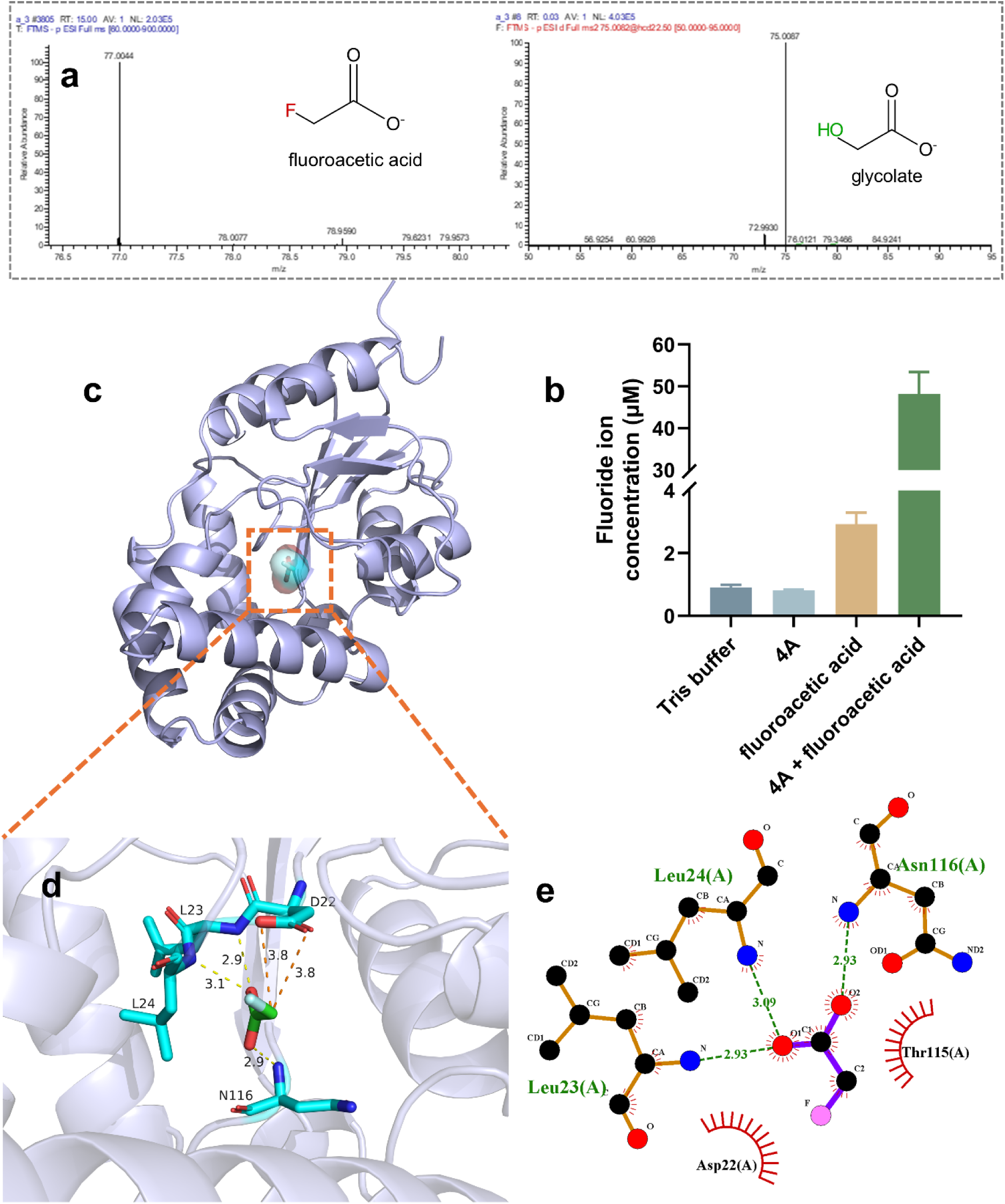
Biochemical and structural evidence for 4A-mediated defluorination of fluoroacetic acid. (a) Representative mass spectrum confirming the formation of glycolate following incubation of purified 4A (10 µM) with fluoroacetic acid (1 mM). (b) Fluoride ion concentrations measured in scaled-up reactions, demonstrating enzyme-dependent fluoride release. Data represent the mean ± standard deviation of two independent biological replicates. (c) Overview of the docking model showing fluoroacetic acid positioned within the active-site cavity of 4A. (d) Enlarged three-dimensional view of the active site highlighting key hydrogen-bonding and nonpolar interactions between fluoroacetic acid and surrounding residues. Hydrogen-bonding interactions are indicated by yellow dashed lines, whereas orange dashed lines denote key interatomic distances between Asp22 and the substrate α-carbon. Amino acid residues are labeled using single-letter codes (N = Asn, L = Leu, T = Thr, and D = Asp). (e) Two-dimensional interaction map summarizing substrate–residue contacts and the proximity of the substrate α-carbon to the catalytic Asp22.

To gain structural insight into fluoroacetic acid recognition by HAD 4A, its X-ray structure was obtained at 1.59 Å resolution (Table S2). The structure of 4A resembled that of other HAD-like dehalogenases, with a core β-sheet comprised of parallel β-strands sandwiched between two α-helix rich regions.^35^ The structure for the monomers of 4A was superposed with two previously solved structures of HAD-like dehalogenases, RJO0230 and POL0530 (PDB IDs 3UMG and 3UM9), yielding backbone RMSD values of 1.30 Å for RJO0230 versus 4A (89 pruned atom pairs) and 1.16 Å for POL0530 versus 4A (119 pruned atom pairs) (Figure 2). One of the major differences in structure for the three HAD-like proteins is the loop between β-strands 5 and 6 (residues 189–205 in 4A), which forms a lid over the active site. In the 4A monomer, this lid covers less of the active site, leaving it more open to solvent than in RJO0230 and POL0530. The relative positions of α-helices 3 and 10 in 4A, which also contribute to the entrance to the active site, are more distant than for the same helices in RJO0230 and POL0530 and further increase the solvent accessibility of the 4A active site compared with those on the other two HAD-like dehalogenases.

**Figure 2.**
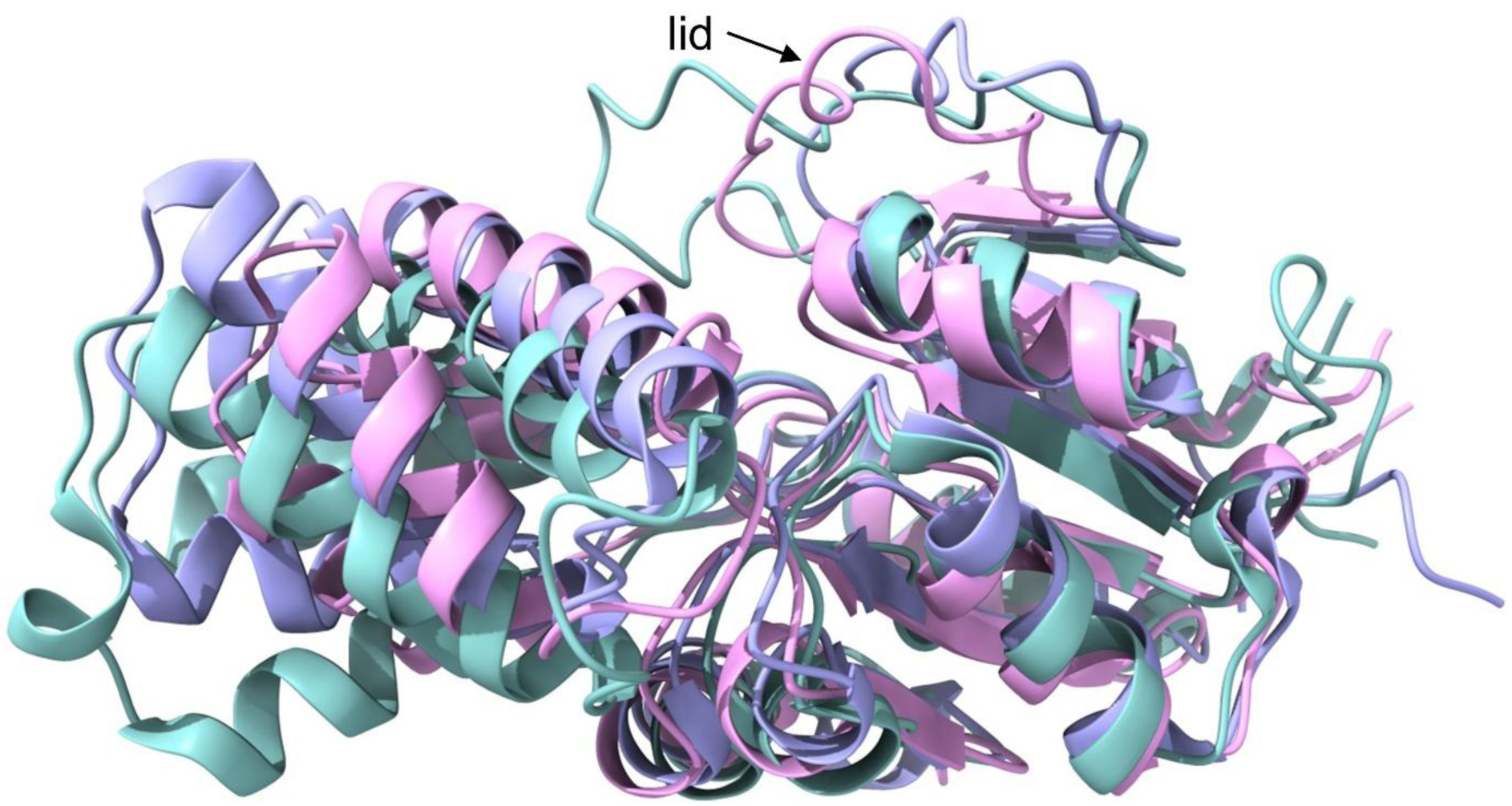
Structural superposition of monomeric HAD-like dehalogenases. The structures of 4A (pink, PDB ID: 9C9E), RJO0230 (teal, PDB ID: 3UMG), and POL0530 (pale lavender blue, PDB ID: 3UM9) were superimposed using the MatchMaker tool in ChimeraX.

Molecular docking simulations for fluoroacetic acid and HAD 4A were performed. The docking model places fluoroacetic acid within the active-site cavity of HAD 4A, with the substrate oriented toward the conserved catalytic center (Figure 1c). Examination of the active site showed that the substrate carboxylic acid was stabilized by hydrogen-bonding interactions with the backbone nitrogens of Leu23, Leu24, and Asn116, with additional nonpolar contacts involving Thr115 (Figure 1d and 1e). Importantly, the α-carbon of fluoroacetic acid was positioned approximately 3.8 Å from the side-chain carboxylate of the catalytic Asp22 (Figure 1d), within a distance range commonly associated with nucleophilic attack in HAD-type dehalogenase-like enzymes.^11,36^

### Delftia acidovorans defluorinase 4A exhibits promiscuous defluorination activity toward PFOA

Given the suggested link between a close homologue of HAD 4A and potential PFOA metabolism, we performed molecular docking analysis, which showed that PFOA could be accommodated within the enzyme’s active-site pocket (Figure 3a). No successful docking was possible with the structures of POL0530 and RJO0230. A systematic screening study that included RJO0230 and POL0530 found that neither enzyme showed detectable defluorination activity toward PFOA, consistent with the absence of successful binding poses found here.^10^

**Figure 3.**
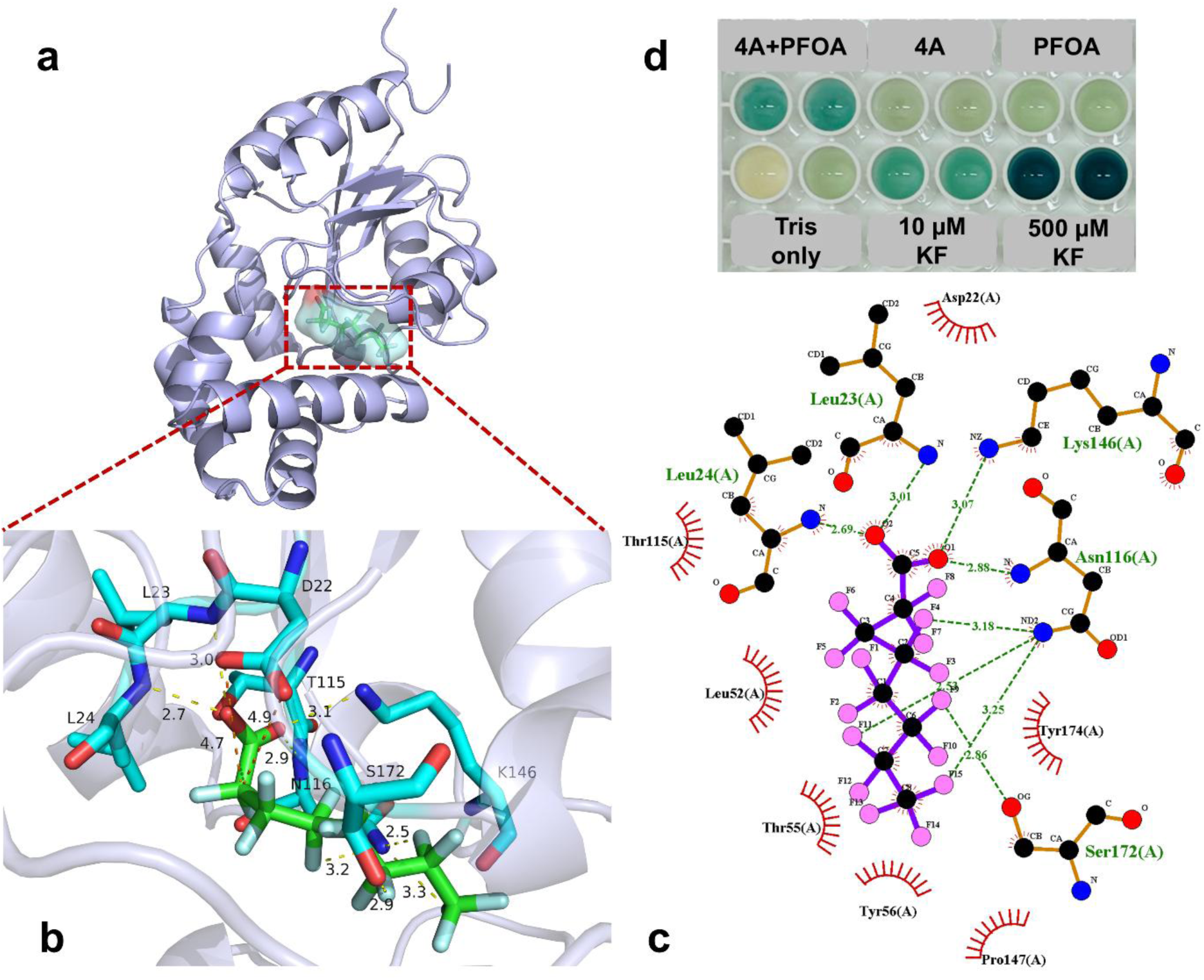
Docking-predicted binding of PFOA to 4A and biosensor-based screening of defluorination activity. (a) Docking model showing PFOA bound within the active-site pocket of 4A. (b) Enlarged view of the PFOA-binding site, illustrating predicted hydrogen-bonding interactions between the PFOA carboxylate group and surrounding active-site residues. (c) Two-dimensional interaction map of the PFOA–4A complex, highlighting hydrogen-bonding interactions involving the carboxylate group (green dashed lines) and hydrophobic or polar contacts between PFOA and active-site residues. (d) Biosensor-based plate assay detecting fluoride release following high-concentration, scaled-up reactions. PFOA (0.5 mM) was incubated with purified 4A (500 µM) at 20℃ for 120 h prior to analysis. Following incubation, aliquots from the reaction mixture were analyzed using a fluoride-responsive riboswitch biosensor with 5-bromo-4-chloro-3-indolyl-β-D-galactopyranoside (X-gal) as the chromogenic substrate. Enzyme-only, substrate-only, buffer-only controls, and fluoride standards (10 and 500 μM KF) are shown.

The lowest-energy docking pose for HAD 4A positioned the PFOA molecule along the substrate-binding channel, with its carboxylate head group oriented toward the catalytic center, while the perfluorinated alkyl chain extended into a hydrophobic cavity. A closer examination of the docked complex showed that the carboxylate group of PFOA was positioned within hydrogen-bonding distance of several residues involved in substrate recognition, including Leu23, Leu24, Asn116, and Lys146 (Figure 3b and 3c). This binding geometry differs from that observed for fluoroacetic acid, indicating a distinct coordination pattern for the longer-chain fluorinated substrate. The catalytically important residue Asp22 was positioned near the α-carbon of PFOA, although at a greater distance (4.7–4.9 Å) than that observed for fluoroacetic acid (3.8 Å) (Figure 3b).

Given the docking-predicted accommodation of PFOA within the 4A active site (Figure 3a–3c), we next tested whether fluoride release could be detected. As this reaction was likely non-physiological and therefore expected to have a low catalytic rate, we tested it with high enzyme (500 µM) and substrate (0.5 mM) loading and prolonged incubation (120 hr). Following incubation, aliquots from the reaction mixture were withdrawn and analyzed using the fluoride-responsive riboswitch biosensor. A clear blue coloration was observed only in biosensor assays of samples derived from reactions containing both PFOA and 4A, whereas enzyme-only and substrate-only controls remained unchanged (Figure 3d), indicating fluoride release after prolonged co-incubation at high concentration.

Quantitative measurement of fluoride ions released during the scaled-up reactions further confirmed these observations (Figure 4). Incubation of PFOA (0.5 mM) with HAD 4A (10 µM and 500 µM) resulted in significant fluoride release at the higher enzyme loading compared with the controls (PFOA alone, HAD 4A alone, or buffer alone). Fluoride release from the 500 µM HAD 4A loading was significantly greater than that observed for the 10 µM HAD 4A incubation. As expected, incubation of fluoroacetic acid (0.5 mM) with HAD 4A (500 µM) yielded substantially higher fluoride release, serving as a positive control and confirming the retained catalytic activity of the enzyme under identical conditions. Together, these results indicate that detectable fluoride release from PFOA was observed only under conditions of high enzyme loading and prolonged incubation.

**Figure 4.**
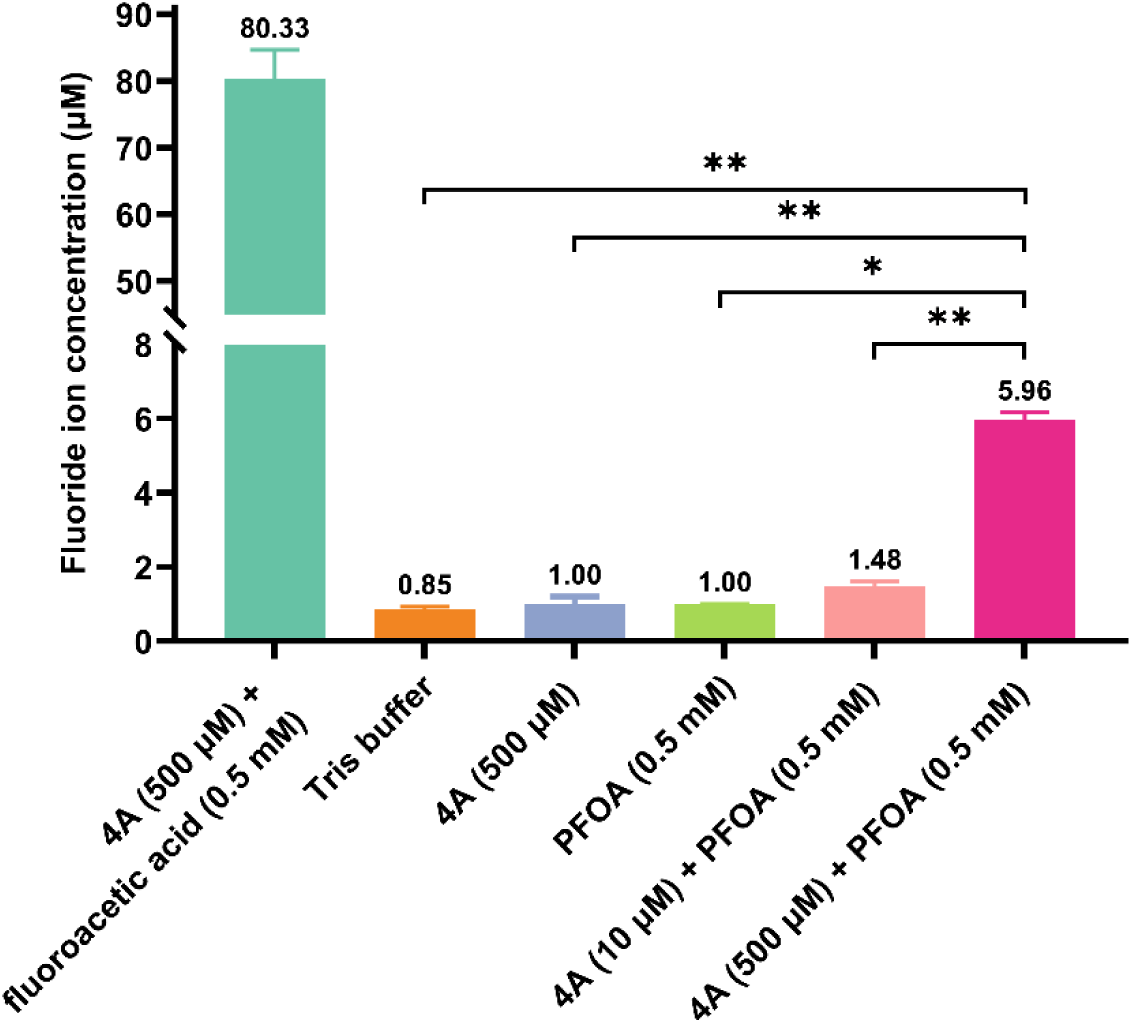
Quantification of fluoride release from PFOA incubated with purified 4A. PFOA (0.5 mM) was incubated with purified 4A at two enzyme loadings (10 and 500 µM) at 20℃ for 120 h. Negative controls included buffer only, 4A only (500 µM), and PFOA only (0.5 mM). A positive control reaction containing fluoroacetic acid (0.5 mM) and 4A (500 µM) was included to confirm enzymatic activity under identical conditions. Fluoride concentrations were quantified using a fluoride ion-selective electrode. Error bars represent the standard deviation of technical measurements obtained from triplicate reactions. Statistical significance was assessed using two-tailed unpaired Welch’s *t*-tests. *, *P* < 0.05; **, *P* < 0.01.

### HAD 4A defluorinates branched PFOA isomers

High-resolution mass spectrometry (HRMS) was used to further characterize the defluorination products formed during incubation of PFOA (0.5 mM) with a high concentration of purified 4A (500 µM). As shown in Figure 4, only a low level of fluoride release (5.9 µM) was detected. While this could simply reflect the low catalytic rate of the reaction, it could also indicate that a minor organofluoride contaminant of the PFOA was the ‘true’ substrate, especially given that PFAS compounds are often contaminated by geometric isomers of the major PFAS product.^37^ HRMS analysis revealed that PFOA stock solution used in this study consisted predominantly of linear PFOA (∼98%), with branched isomers accounting for approximately 1.92% of the total PFOA composition (Figure 5).^38^

**Figure 5.**
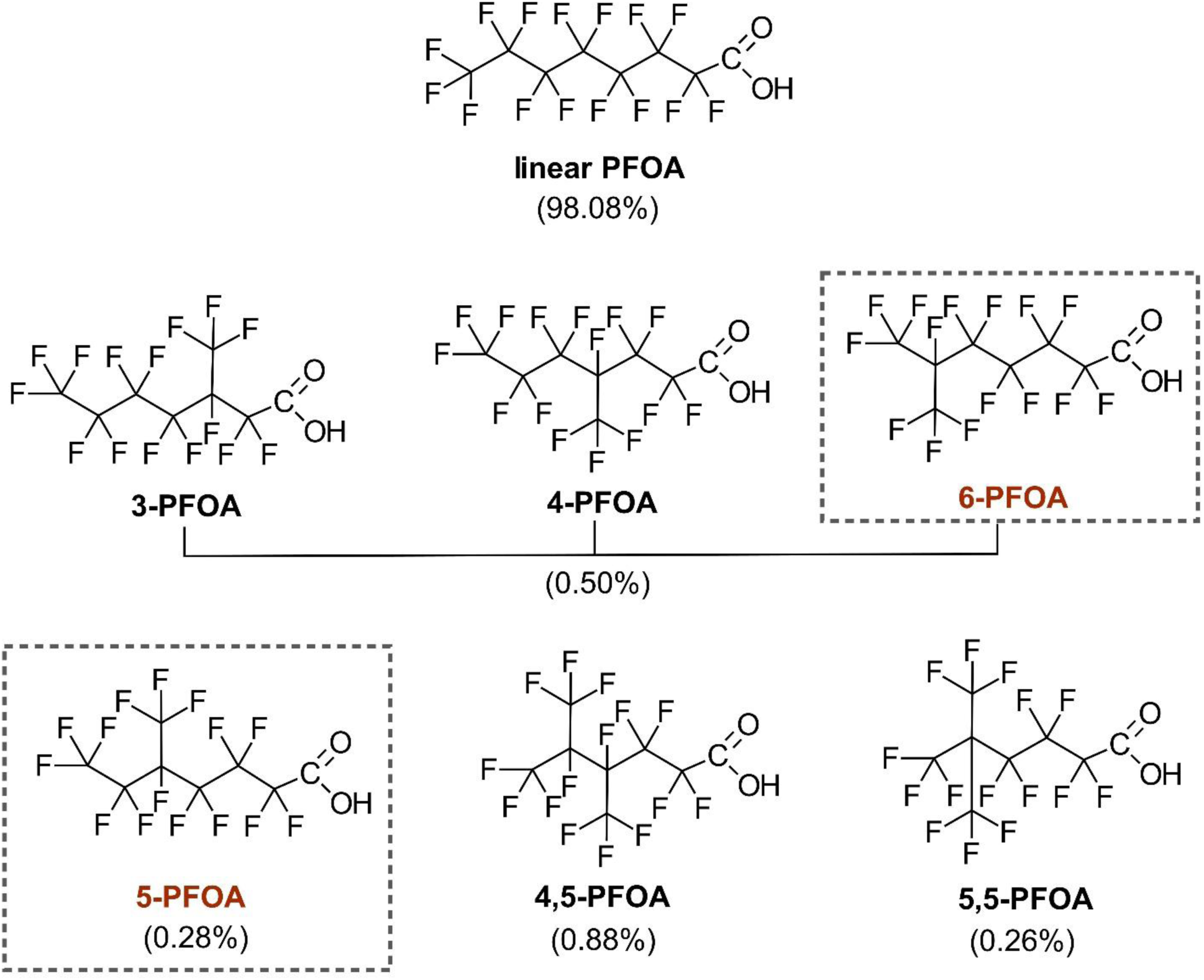
Chemical structures of linear and branched PFOA isomers present in the PFOA stock solution used in this study. Values in parentheses indicate the relative abundance (%) of each isomer as reported by Sun et al.^38^. Dashed outlines indicate branched PFOA isomers associated with detectable fluoride release in the presence of HAD 4A under the conditions tested. Abbreviations: 3-PFOA, perfluoro-3-methylheptanoic acid; 4-PFOA, perfluoro-4-methylheptanoic acid; 5-PFOA, perfluoro-5-methylheptanoic acid; 6-PFOA, perfluoro-6-methylheptanoic acid; 4,5-PFOA, perfluoro-4,5-dimethylhexanoic acid; 5,5-PFOA, perfluoro-5,5-dimethylhexanoic acid.

HRMS analysis of the enzymatic reaction sample revealed the formation of a distinct product ion at *m/z* 407, which was absent from all control reactions lacking enzyme or substrate (Figure 6a). The observed *m/z* value matches the theoretical mass of a defluorinated PFOA-derived product and may correspond to either perfluoro-3,4,5-trihydroxy-6-methyloctanoic acid or perfluoro-3,4,5-trihydroxy-5-methyloctanoic acid (theoretical *m/z* = 407), both of which are predicted products of branched PFOA defluorination. Tandem MS (MS/MS) fragmentation analysis of the m/z 407 ion revealed characteristic fragment ions at *m/z* 175, 219, 238, and 275 (Figures S3 and S4). These fragments closely matched the predicted fragmentation patterns of defluorination products derived from 5-PFOA or 6-PFOA. Notably, the fragment at *m/z* 219 has been reported as a characteristic ion associated with 5-PFOA- and 5,5-PFOA-derived structures,^39^ providing additional support for selective enzymatic transformation of specific branched PFOA isomers. However, due to overlapping fragmentation pathways among branched isomers, the precise identity of the enzymatically transformed isomer cannot be unambiguously assigned based solely on MS/MS data.

**Figure 6.**
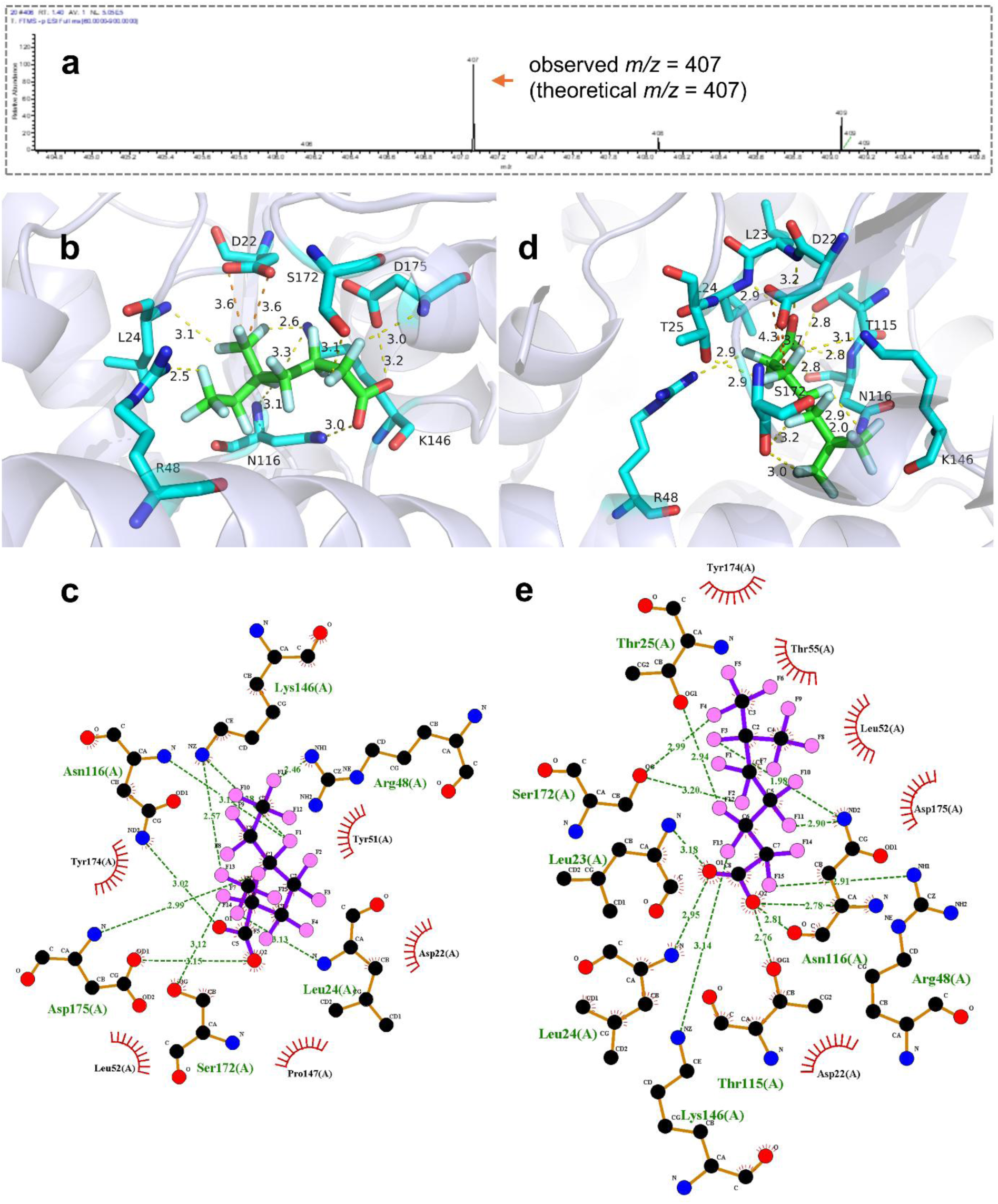
HRMS identification and isomer-specific docking analysis of PFOA defluorination products formed by 4A. (a) High-resolution mass spectrum of the enzymatic reaction sample (0.5 mM PFOA and 500 µM 4A) showing a product ion at *m/z* 407, consistent with the theoretical mass of a defluorinated PFOA-derived product. (b, c) Predicted binding pose (b) and corresponding two-dimensional interaction map (c) of 5-PFOA within the active site of 4A. (d, e) Predicted binding pose (d) and corresponding two-dimensional interaction map (e) of 6-PFOA within the active site of 4A.

Molecular docking was subsequently performed using 5-PFOA and 6-PFOA to evaluate their binding within the active site of HAD 4A. Docking analysis showed that both isomers could be accommodated within the same active-site cavity of HAD 4A, although they adopted distinct binding conformations (Figure 6b–6e). For 5-PFOA, the carboxylate group is stabilized primarily through hydrogen-bonding interactions involving Asn116 and Asp175 (Figure 6c). In contrast, the carboxylate of 6-PFOA engages a different polar network featuring Asn116 and Thr115, together with additional backbone-associated polar contacts in the vicinity of Leu23 and Leu24 (Figure 6e).

To maintain consistency with the docking analyses performed for fluoroacetic acid and linear PFOA, the spatial relationship between the catalytic Asp22 and the substrate backbone was also examined for the branched PFOA isomers. In the 5-PFOA docking model, Asp22 is positioned approximately 3.6 Å from the substrate backbone carbon proximal to the branching region (Figure 6b). In the 6-PFOA model, the corresponding distance ranges from ∼3.7 to 4.3 Å (Figure 6d), reflecting a slightly altered substrate orientation within the active site.

Despite these conserved geometric features, the two isomers exhibited distinct local interaction patterns within the hydrophobic region of the binding pocket, as evidenced by differences in residue contacts and substrate orientation in the docking models for 5-PFOA and 6-PFOA.

## DISCUSSION

Previous work reported that a slight increase in fluoride release for a HAD from *D*. *acidovorans* under whole-cell conditions (PZP66635.1).^18^ We had unsuccessfully attempted to express and purify PZP66635.1 for biochemical and structural studies. Instead, we focused on HAD 4A, a close homolog of PZP66635.1 from *D*. *acidovorans* strain D4B,^19^ which was amenable to purification and crystallization, enabling direct structural analysis of defluorination-relevant activity.

We have shown that purified HAD 4A catalyzed defluorination of fluoroacetic acid, demonstrating that, although not annotated as a fluoroacetate dehalogenase, it is one of only a few HADs reported to cleave C–F bonds.^9^ Surprisingly, we also found that HAD 4A can defluorinate a perfluorinated compound, specifically the geometric isomers of PFOA. To the best of our knowledge, this is the first demonstration of a specific hydrolase that defluorinates perfluorinated compounds.

Structural analysis suggests that the substrate-binding pocket of HAD 4A is both larger and more open to solvent than those of other HADs (specifically POL0530 and RJO0230), allowing it to accommodate large perfluorinated compounds. Although this insight was obtained using linear PFOA as the ligand in docking studies, the catalytic aspartate appears too distant from a fluorinated carbon to catalyze detectable defluorination of linear PFOA. Instead, the geometric isomers 5-PFOA and 6-PFOA were shown to be substrates for HAD 4A, with docking suggesting that these isomers position the fluorinated carbons within range of S_N_2 nucleophilic attack by the conserved catalytic aspartic acid residue.

While we were conducting our investigation, Farajollahi et al. published a study in which they were unable to detect PFOA defluorinase activity in HAD 4A (termed DeHa5 in their study).^40^ While this seems to contradict our findings, there are at least two explanations that may account for the discrepancy. Farajollahi et al. used lower enzyme (∼20.7 µM) and substrate (1 mM) concentrations, which may have been too low to detect HAD 4A’s promiscuous PFAS defluorinase activity. Alternatively, the composition of the PFOA solution used in their investigation may have contained lower concentrations of 5-PFOA and 6-PFOA, the actual substrates for HAD 4A. This also highlights the lack of information about the composition of available PFAS compounds, which are often uncharacterized ensembles of isomers, rather than a single compound.

Incubation of PFOA (0.5 mM) with 4A at a high enzyme loading (500 μM) for 120 h yielded only ∼4.96 μM fluoride, corresponding to an apparent minimum turnover rate of 8.3 × 10^−5^ h^−1^ (2 × 10^−8^ s^−1^). The true turnover rate is likely higher than this estimate, given that the substrates were a low-concentration contaminant of linear PFOA, and this value was estimated from an endpoint assay. However, the PFAS-defluorination activity of HAD 4A is likely to proceed at a rate orders of magnitude lower than those reported for known fluoroacetate-degrading enzymes and far below typical physiological enzymatic rates (∼10 s^−1^).^41^ Such low catalytic efficiency suggests that the branched PFOA isomers are not physiological substrates for HAD 4A. Instead, the observed activity is best described as a promiscuous defluorination reaction, defined as a low-level secondary activity distinct from an enzyme’s physiological function.^42^ Although enzymatic promiscuity is generally considered physiologically irrelevant, it can be mechanistically informative and reflects an enzyme’s latent evolutionary potential.^42,43^ In this context, the reproducible detection of fluoride release from PFOA provides quantitative evidence that the HAD-like enzymes defluorinate PFAS via a classical S_N_2 mechanism, albeit inefficiently.

Promiscuous activities provide the biochemical potential upon which natural selection acts to generate new metabolic enzymes and pathways. Confirmation that promiscuous PFAS defluorinase activity is present in the environment raises the question of why PFAS catabolic pathways have not yet evolved. We, and others, have speculated that there must be evolutionary (rather than biochemical) barriers to the evolution of such pathways, such as the intrinsic toxicity of fluoride released during hydrolysis.^9,44^

Although HAD 4A has only promiscuous PFAS activity, it may be a useful scaffold for enzyme engineering. Directed evolution, rational design, or emerging AI protein design strategies may enhance defluorination activity toward fluorinated substrates that are otherwise resistant to biological defluorination. Although the complete mineralization of PFAS remains a formidable long-term challenge, the present findings help define a tractable path forward. The identification of experimentally validated defluorination activity via an SN_2_ mechanism provides a foundation for developing tailored biocatalysts that enhance PFAS defluorination, even if such capabilities remain beyond those of naturally evolved enzymatic systems.

## ASSOCIATED CONTENT

### Supporting Information

The following file is available free of charge.

Tables summarizing 4A sequence information and crystal structure refinement statistics; figures showing protein expression and purification, as well as LC-MS identification of defluorination products from fluoroacetate and branched PFOA isomers (PDF).

## AUTHOR INFORMATION

### Funding Sources

This study was supported by a CSIRO CERC Postdoctoral Fellowship (1310-CERC Fellow). This research was undertaken in part using the MX2 beamline at the Australian Synchrotron, part of ANSTO, and made use of the Australian Cancer Research Foundation (ACRF) detector.

### Notes

The authors declare that they have no known competing financial interests.

## Notes

### Competing Interest Statement

The authors have declared no competing interest.

## REFERENCES

(1) Buck, R. C.; Franklin, J.; Berger, U.; Conder, J. M.; Cousins, I. T.; De Voogt, P.; Jensen, A. A.; Kannan, K.; Mabury, S. A.; van Leeuwen, S. P. Perfluoroalkyl and polyfluoroalkyl substances in the environment: terminology, classification, and origins. Integr. Environ. Assess. Manag. 2011, 7 (4), 513–541.

(2) Glüge, J.; Scheringer, M.; Cousins, I. T.; DeWitt, J. C.; Goldenman, G.; Herzke, D.; Lohmann, R.; Ng, C. A.; Trier, X.; Wang, Z. An overview of the uses of per- and polyfluoroalkyl substances (PFAS). Environ. Sci.: Processes Impacts 2020, 22 (12), 2345–2373.

(3) Johnson, G. R.; Brusseau, M. L.; Carroll, K. C.; Tick, G. R.; Duncan, C. M. Global distributions, source-type dependencies, and concentration ranges of per-and polyfluoroalkyl substances in groundwater. Sci. Total Environ. 2022, 841, 156602.

(4) O’Hagan, D. Understanding organofluorine chemistry. An introduction to the C–F bond. Chem. Soc. Rev. 2008, 37 (2), 308–319.

(5) Wang, Z.; DeWitt, J. C.; Higgins, C. P.; Cousins, I. T. A never-ending story of per- and polyfluoroalkyl substances (PFASs)?. Environm. Sci. Technol. 2017, 51 (5), 2508–2518.

(6) Zhang, Z.; Sarkar, D.; Biswas, J. K.; Datta, R. Biodegradation of per-and polyfluoroalkyl substances (PFAS): A review. Bioresour. Technol. 2022, 344, 126223.

(7) LaFond, J. A.; Hatzinger, P. B.; Guelfo, J. L.; Millerick, K.; Jackson, W. A. Bacterial transformation of per- and poly-fluoroalkyl substances: a review for the field of bioremediation. Environ. Sci.: Adv. 2023, 2 (8), 1019–1041.

(8) Berhanu, A.; Mutanda, I.; Taolin, J.; Qaria, M. A.; Yang, B.; Zhu, D. A review of microbial degradation of per-and polyfluoroalkyl substances (PFAS): Biotransformation routes and enzymes. Sci. Total Environ. 2023, 859, 160010.

(9) Hu, M.; Scott, C. Toward the development of a molecular toolkit for the microbial remediation of per-and polyfluoroalkyl substances. Appl. Environ. Microbiol. 2024, 90 (4), e00157–00124.

(10) Khusnutdinova, A. N.; Batyrova, K. A.; Brown, G.; Fedorchuk, T.; Chai, Y. S.; Skarina, T.; Flick, R.; Petit, A.-P.; Savchenko, A.; Stogios, P.; Yakunin, A. F. Structural insights into hydrolytic defluorination of difluoroacetate by microbial fluoroacetate dehalogenases. FEBS J. 2023, 290 (20), 4966–4983.

(11) Chan, P. W. Y.; Yakunin, A. F.; Edwards, E. A.; Pai, E. F. Mapping the reaction coordinates of enzymatic defluorination. J. Am. Chem. Soc. 2011,133 (19), 7461–7468.

(12) Li, Y.; Yue, Y.; Zhang, H.; Yang, Z.; Wang, H.; Tian, S.; Wang, J.-b.; Zhang, Q.; Wang, W. Harnessing fluoroacetate dehalogenase for defluorination of fluorocarboxylic acids: *in silico* and *in vitro* approach. Environ. Int. 2019, 131, 104999.

(13) Bhardwaj, S.; Hu, M.; Scott, C.; Manefield, M. J. Role of acyl-CoA synthetase (ACS) from *Dietzia aurantiaca* strain J3 and glutathione (GSH) in fluorotelomer biotransformation. Environ. Sci. Technol. 2026, 10.1021/acs.est.5c11041.

(14) Mothersole, R. G.; Wynne, F. T.; Rota, G.; Mothersole, M. K.; Liu, J.; Van Hamme, J. D. Formation of CoA adducts of short-chain fluorinated carboxylates catalyzed by acyl-CoA synthetase from *Gordonia* sp. strain NB4-1Y. ACS Omega 2023, 8 (42), 39437–39446.

(15) Mothersole, R. G.; Mothersole, M. K.; Goddard, H. G.; Liu, J.; Van Hamme, J. D. Enzyme catalyzed formation of CoA adducts of fluorinated hexanoic acid analogues using a long-chain acyl-CoA synthetase from *Gordonia* sp. strain NB4-1Y. Biochemistry 2024, 63 (17), 2153–2165.

(16) Chetverikov, S.; Hkudaygulov, G.; Sharipov, D.; Starikov, S.; Chetverikova, D. Biodegradation potential of C7-C10 perfluorocarboxylic acids and data from the genome of a new strain of *Pseudomonas mosselii* 5(3). Toxics 2023, 11 (12), 1001.

(17) Sorn, S.; Matsuura, N.; Honda, R. Metagenome-assembled genomes and metatranscriptome analysis of perfluorooctane sulfonate-reducing bacteria enriched from activated sludge. Environ. Microbiol. 2025, 27 (4), e70087.

(18) Harris, J. D.; Coon, C. M.; Doherty, M. E.; McHugh, E. A.; Warner, M. C.; Walters, C. L.; Orahood, O. M.; Loesch, A. E.; Hatfield, D. C.; Sitko, J. C. Engineering and characterization of dehalogenase enzymes from *Delftia acidovorans* in bioremediation of perfluorinated compounds. Synth. Syst. Biotechnol. 2022, 7 (2), 671–676.

(19) Harris, J.; Gross, M.; Kemball, J.; Farajollahi, S.; Dennis, P.; Sitko, J.; Steel, J. J.; Almand, E.; Kelley-Loughnane, N.; Varaljay, V. A. Draft genome sequence of the bacterium *Delftia acidovorans* strain D4B, isolated from soil. Microbiol. Resour. Announc. 2021, 10, 10.1128/mra.00635-00621.

(20) Hu, M.; Bhardwaj, S.; Martinez, J. P. O.; Manefield, M. J.; Scott, C. A fluoride-specific riboswitch biosensor for high-throughput enzymatic defluorination screening. ACS Synth. Biol. 2025, 14 (11), 4322–4329.

(21) Abrahams, G.; Newman, J. Data- and diversity-driven development of a Shotgun crystallization screen using the Protein Data Bank. Acta Crystallogr. D Struct. Biol. 2021, 77 (11), 1437–1450.

(22) Aragão, D.; Aishima, J.; Cherukuvada, H.; Clarken, R.; Clift, M.; Cowieson, N. P.; Ericsson, D. J.; Gee, C. L.; Macedo, S.; Mudie, N.; Panjikar, S.; Price, J. R.; Riboldi-Tunnicliffe, A.; Rostan, R.; Williamson, R.; Caradoc-Davies, T. T. MX2: a high-flux undulator microfocus beamline serving both the chemical and macromolecular crystallography communities at the Australian Synchrotron. J. Synchrotron Rad. 2018, 25 (3), 885–891.

(23) Vonrhein, C.; Flensburg, C.; Keller, P.; Sharff, A.; Smart, O.; Paciorek, W.; Womack, T.; Bricogne, G. Data processing and analysis with the autoPROC toolbox. Acta Crystallogr. D Struct. Biol. 2011, 67 (4), 293–302.

(24) Kabsch, W. XDS. Acta Crystallogr. D Struct. Biol. 2010, 66 (2), 125–132.

(25) Agirre, J.; Atanasova, M.; Bagdonas, H.; Ballard, C. B.; Basle, A.; Beilsten-Edmands, J.; Borges, R. J.; Brown, D. G.; Burgos-Marmol, J. J.; Berrisford, J. M.; Bond, P. S.;, et al. The CCP4 suite: integrative software for macromolecular crystallography. Acta Crystallogr. D Struct. Biol. 2023, 79 (6), 449–461.

(26) French, S.; Wilson, K. On the treatment of negative intensity observations. Acta Crystallogr. 1978, 34 (4), 517–525.

(27) McCoy, A. J.; Grosse-Kunstleve, R. W.; Adams, P. D.; Winn, M. D.; Storoni, L. C.; Read, R. J. *Phaser* crystallographic software. J. Appl. Crystallogr. 2007, 40 (4), 658–674.

(28) Jumper, J.; Evans, R.; Pritzel, A.; Green, T.; Figurnov, M.; Ronneberger, O.; Tunyasuvunakool, K.; Bates, R.; Žídek, A.; Potapenko, A.; Bridgland, A.;, et al. Highly accurate protein structure prediction with AlphaFold. Nature 2021, 596 (7873), 583–589.

(29) Emsley, P.; Lohkamp, B.; Scott, W. G.; Cowtan, K. Features and development of *Coot*. Acta Crystallogr. D Struct. Biol. 2010, 66 (4), 486–501.

(30) Bricogne, G.; B. E.; Brandl, M.; Flensburg, C.; Keller, P.; Paciorek, W.; Roversi, P.; Sharff, A.; Smart, O. S.; Vonrhein, C.; Womack, T. O. BUSTER, Global Phasing Ltd., Cambridge, United Kingdom, 2017.

(31) Liebschner, D.; Afonine, P. V.; Baker, M. L.; Bunkóczi, G.; Chen, V. B.; Croll, T. I.; Hintze, B.; Hung, L.-W.; Jain, S.; McCoy, A. J. Macromolecular structure determination using X-rays, neutrons and electrons: recent developments in Phenix. Acta Crystallogr., Sect. D: Struct. Biol. 2019, 75 (10), 861–877.

(32) Eberhardt, J.; Santos-Martins, D.; Tillack, A. F.; Forli, S. AutoDock Vina 1.2.0: New docking methods, expanded force field, and Python bindings. J. Chem. Inf. Model. 2021, 61 (8), 3891–3898.

(33) Schrödinger LLC. The PyMOL Molecular Graphics System, Version 3.0, Schrödinger LLC, 2024.

(34) Laskowski, R. A.; Swindells, M. B. LigPlot+: Multiple ligand–protein interaction diagrams for drug discovery. J. Chem. Inf. Model. 2011, 51 (10), 2778–2786.

(35) Chan, P. W.; Chakrabarti, N.; Ing, C.; Halgas, O.; To, T. K.; Wälti, M.; Petit, A. P.; Tran, C.; Savchenko, A.; Yakunin, A. F. Defluorination capability of l-2-haloacid dehalogenases in the HAD-like hydrolase superfamily correlates with active site compactness. ChemBioChem 2022, 23 (1), e202100414.

(36) Yue, Y.; Fan, J.; Xin, G.; Huang, Q.; Wang, J.-b.; Li, Y.; Zhang, Q.; Wang, W. Comprehensive understanding of fluoroacetate dehalogenase-catalyzed degradation of fluorocarboxylic acids: A QM/MM approach. Environ. Sci. Technol. 2021, 55 (14), 9817–9825.

(37) Londhe, K.; Lee, C. S.; McDonough, C. A.; Venkatesan, A. K. The need for testing isomer profiles of perfluoroalkyl substances to evaluate treatment processes. Environ. Sci. Technol. 2022, 56 (22), 15207–15219.

(38) Sun, J.; Yu, T. T.; Mirabediny, M.; Lee, M.; Jones, A.; O’Carroll, D. M.; Manefield, M. J.; Kumar, P. V.; Pickford, R.; Ramadhan, Z. R.; Bhattacharyya, S. K.; Åkermark, B.; Das, B.; Kumar, N. Soluble metal porphyrins-Zero-valent zinc system for effective reductive defluorination of branched per and polyfluoroalkyl substances (PFASs). Water Res. 2024, 258, 121803.

(39) Pellizzaro, A.; Zaggia, A.; Fant, M.; Conte, L.; Falletti, L. Identification and quantification of linear and branched isomers of perfluorooctanoic and perfluorooctane sulfonic acids in contaminated groundwater in the veneto region. J. Chromatogr. A 2018, 1533, 143–154.

(40) Farajollahi, S.; Lombardo, N. V.; Crenshaw, M. D.; Guo, H. B.; Doherty, M. E.; Davison, T. R.; Steel, J. J.; Almand, E. A.; Varaljay, V. A.; Suei-Hung, C.; Mirau, P. A.; Berry, R. J.; Kelley-Loughnane, N.; Dennis, P. B. Defluorination of organofluorine compounds using dehalogenase enzymes from *Delftia acidovorans* (D4B). ACS Omega 2024, 9 (26), 28546–28555.

(41) Bar-Even, A.; Noor, E.; Savir, Y.; Liebermeister, W.; Davidi, D.; Tawfik, D. S.; Milo, R. The moderately efficient enzyme: Evolutionary and physicochemical trends shaping enzyme parameters. Biochemistry 2011, 50 (21), 4402–4410.

(42) Tawfik, O. K.; S., D. Enzyme promiscuity: A mechanistic and evolutionary perspective. Annu. Rev. Biochem. 2010. 79 (1), 471–505.

(43) Babtie, A.; Tokuriki, N.; Hollfelder, F. What makes an enzyme promiscuous? Curr. Opin. Chem. Biol. 2010, 14 (2), 200–207.

(44) Wackett, L. P. Evolutionary obstacles and not C–F bond strength make PFAS persistent. Microb. Biotechnol. 2024, 17, e14463.

