## Supporting Information for "Selective Hydrolytic Defluorination of Branched Perfluorooctanoic Acid Isomers by a Haloacid Dehalogenase"

[†]Present affiliation.

School of Molecular Sciences, The University of Western Australia, Perth, WA 6009, Australia

 (C.S.)

**Table S1. 4A sequences.**

| **Enzyme name** | **4A** |
|---|---|
| Gene name | 4A |
| NCBI ID | MCA1067727.1 |
| PDB | 9C9E |
| Species | *Delftia acidovorans* strain D4B |
| References | Harris et al.[1,2] |
| Protein sequence<br><br>(224 aa) | MNTPAPLTETTAAAWPRAVLFDLLTALLDSWTVWNSAAGSEAAGRDWRAE<br>YLRLTYGCGAYQPYEDLVREAARNRGLPASAADRLEAQWDQLQPWDGAR<br>ELLAALRPHCRLAVVTNCSERLGQRAAALLGVDWDVVVTSEAAGFYKPDP<br>RPYQLALDRLGLPADQAAFVAGSGYDLFGTSAVGLRTFWHNRVGLSRPAG<br>APAAEGEAATLAPALPWLRGFAAGR |
| DNA sequence before optimization<br><br>(675 bp) | ATGAACACCCCAGCCCCACTGACGGAAACGACCGCCGCCGCCTGGCCCC<br>GTGCCGTGCTTTTCGACCTGCTGACCGCCTTGCTCGACTCCTGGACGGTA<br>TGGAACAGCGCTGCCGGCAGCGAGGCCGCAGGGCGCGACTGGCGCGCCG<br>AGTACCTGCGCCTGACCTATGGCTGCGGCGCCTACCAGCCCTATGAAGAC<br>CTGGTGCGCGAGGCGGCCCGCAACCGGGGCCTGCCCGCCTCGGCCGCCG<br>ACCGGCTGGAGGCGCAATGGGACCAGCTGCAGCCCTGGGATGGCGCACG<br>CGAGCTGCTGGCCGCCCTGCGCCCGCACTGCCGCCTGGCCGTGGTGACCA<br>ATTGCTCCGAGCGCCTGGGTCAACGTGCCGCCGCGCTGCTGGGCGTGGAT<br>TGGGATGTGGTGGTCACCTCGGAAGCAGCGGGCTTCTACAAGCCCGACC<br>CCCGGCCCTACCAGCTGGCGCTGGACCGGCTGGGGCTGCCTGCGGACCA<br>GGCCGCCTTTGTGGCCGGATCGGGCTATGACCTGTTCGGCACCTCGGCCG<br>TGGGCCTGCGCACCTTCTGGCACAACCGCGTGGGCCTGTCGCGGCCTGCG<br>GGCGCGCCTGCTGCCGAGGGGGAAGCCGCCACGCTGGCCCCCGCCTTGC<br>CCTGGCTGCGCGGCTTTGCCGCAGGCCGCTGA |
| DNA sequence after optimization<br><br>(675 bp) | ATGAATACGCCAGCTCCCCTAACAGAAACTACAGCGGCGGCTTGGCCGC<br>GCGCTGTGCTGTTCGACCTGTTGACCGCACTGTTGGACTCCTGGACCGTT<br>TGGAACAGCGCGGCGGGTTCTGAGGCAGCCGGCAGAGATTGGCGTGCAG<br>AATACCTGCGCTTGACGTACGGGTGCGGTGCGTATCAACCGTACGAAGA<br>TCTGGTTCGTGAGGCAGCCCGCAACCGCGGTCTGCCGGCAAGCGCTGCG<br>GACCGCCTTGAGGCACAGTGGGACCAGCTGCAGCCTTGGGACGGCGCAA<br>GAGAACTGTTAGCCGCGTTGCGTCCGCATTGCCGTTTGGCGGTGGTGACC<br>AATTGTAGCGAGCGCCTGGGCCAGCGTGCGGCGGCGTTGCTCGGTGTAG<br>ATTGGGACGTTGTGGTCACTTCCGAGGCGGCGGGCTTCTATAAACCGGAT<br>CCGCGTCCGTACCAACTGGCTCTGGATCGTCTGGGTCTCCCAGCCGACCA<br>AGCGGCGTTTGTTGCAGGCAGCGGTTATGATTTATTTGGTACGTCGGCTG<br>TGGGCCTGCGGACCTTTTGGCACAACCGTGTTGGCCTGAGCCGTCCGGCC<br>GGCGCACCGGCGGCGGAAGGTGAAGCGGCGACCCTGGCGCCAGCTCTGC<br>CGTGGCTGCGTGGTTTCGCCGCGGGTCGTTAA |

**Table S2. Crystal structure processing and refinement statistics.**

| | **4A** |
|---|---|
| **PDB ID** | 9C9E |
| **Data reduction** | |
| Space group | $P2_12_12_1$ |
| Cell dimensions | |
| *a*, *b*, *c* (Å) | 43.2, 68.5, 69.9 |
| α, β, γ (º) | 90.0, 90.0, 90.0 |
| Resolution (Å) | 48.9–1.59 (1.61–1.59) |
| $R_{meas}$ | 0.153 (3.81) |
| $R_{pim}$ | 0.052 (1.75) |
| *I* / *I* | 7.8 (0.4) |
| $CC_{1/2}$ | 0.998 (0.528) |
| Completeness | 94.2 (91.8) |
| Multiplicity | 8.1 (4.5) |
| **Refinement** | |
| Resolution (Å) | 48.93–1.59 |
| No. reflections | 26822 |
| $R_{\mathrm{work}}$ / $R_{\mathrm{free}}$ | 0.2256/0.2487 |
| No. atoms | |
| Protein | 1612 |
| Ligand/ion | 6 |
| Water | 123 |
| *B*-factors | |
| Protein | 26.8 |
| Ligand/ion | 46. |
| Water | 39.9 |
| R.m.s. deviations | |
| Bond lengths (Å) | 1.00 |
| Bond angles (°) | 0.009 |

*Values in parentheses are for highest-resolution shell.

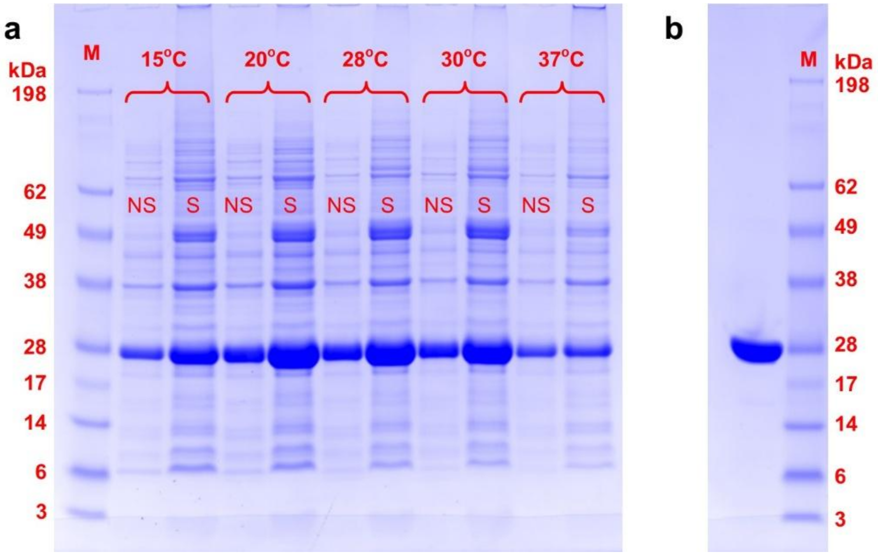

**Figure S1. SDS-PAGE analysis of heterologous expression and purification of 4A.**
(**a**) Expression of 4A at different induction temperatures (15, 20, 28, 30, and 37°C). "NS" and "S" indicate the insoluble and soluble fractions, respectively. For the insoluble fraction, 5 μL of sample was mixed with 15 μL of Milli-Q water and 5 μL of 5× Gel Loading Dye (Thermo Fisher). For the soluble fraction, 20 μL of sample was mixed with 5 μL of 5× Gel Loading Dye. After denaturation at 95°C for 5 min, 10 μL of each mixture was loaded per lane. (**b**) Purified 4A. A total of 20 μL of purified protein (1.5 mg/mL) was loaded. MW marker: SeeBlue Pre-stained Protein Standard (Thermo Fisher).

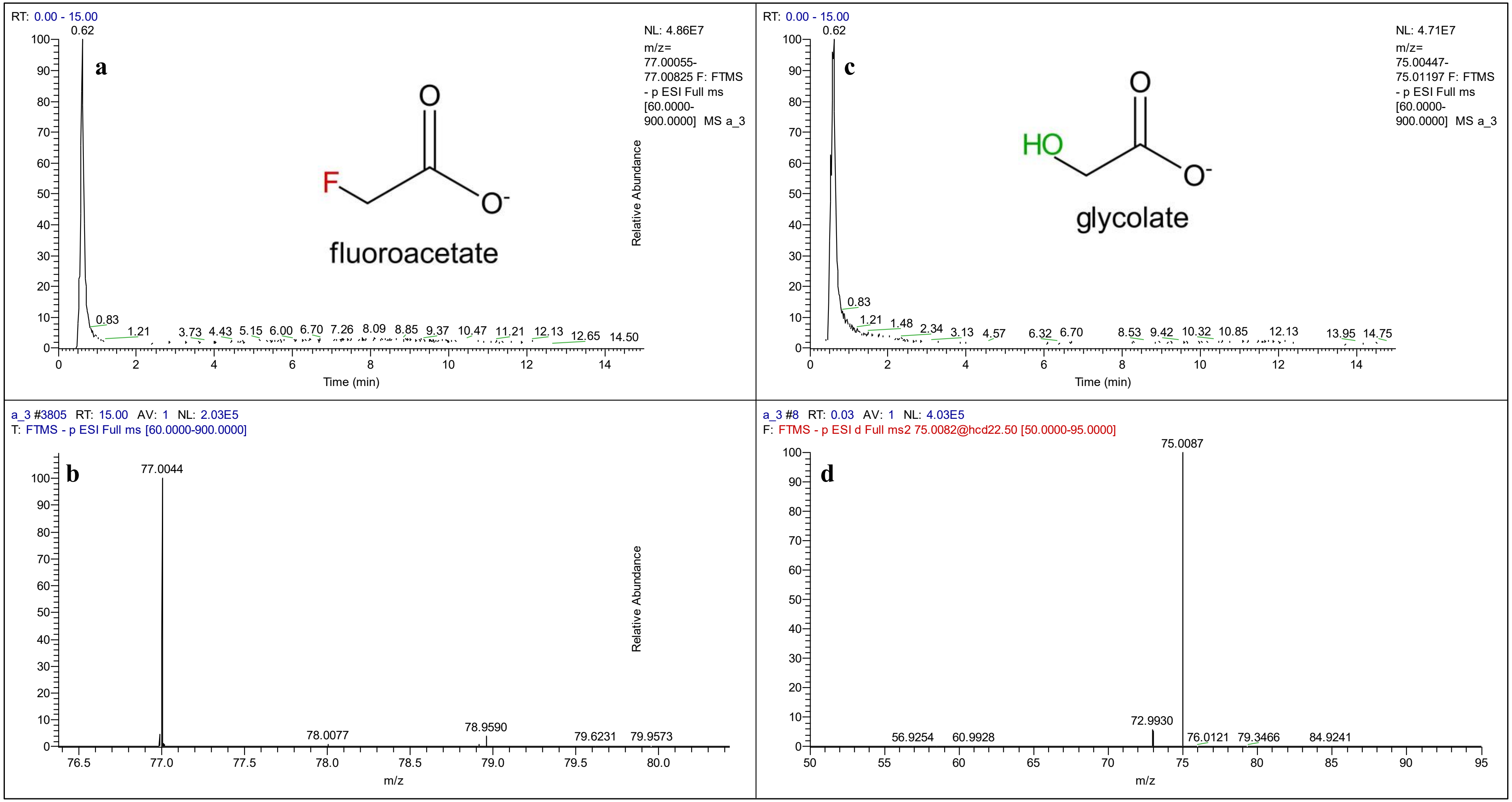

**Figure S2 Defluorination activity of 4A toward fluoroacetate analyzed by LC-MS.**
(**a**, **b**) LC (a) and MS (b) spectra of fluoroacetate. (**c**, **d**) LC (c) and MS (d) spectra of glycolate produced after defluorination of fluoroacetate by 4A.

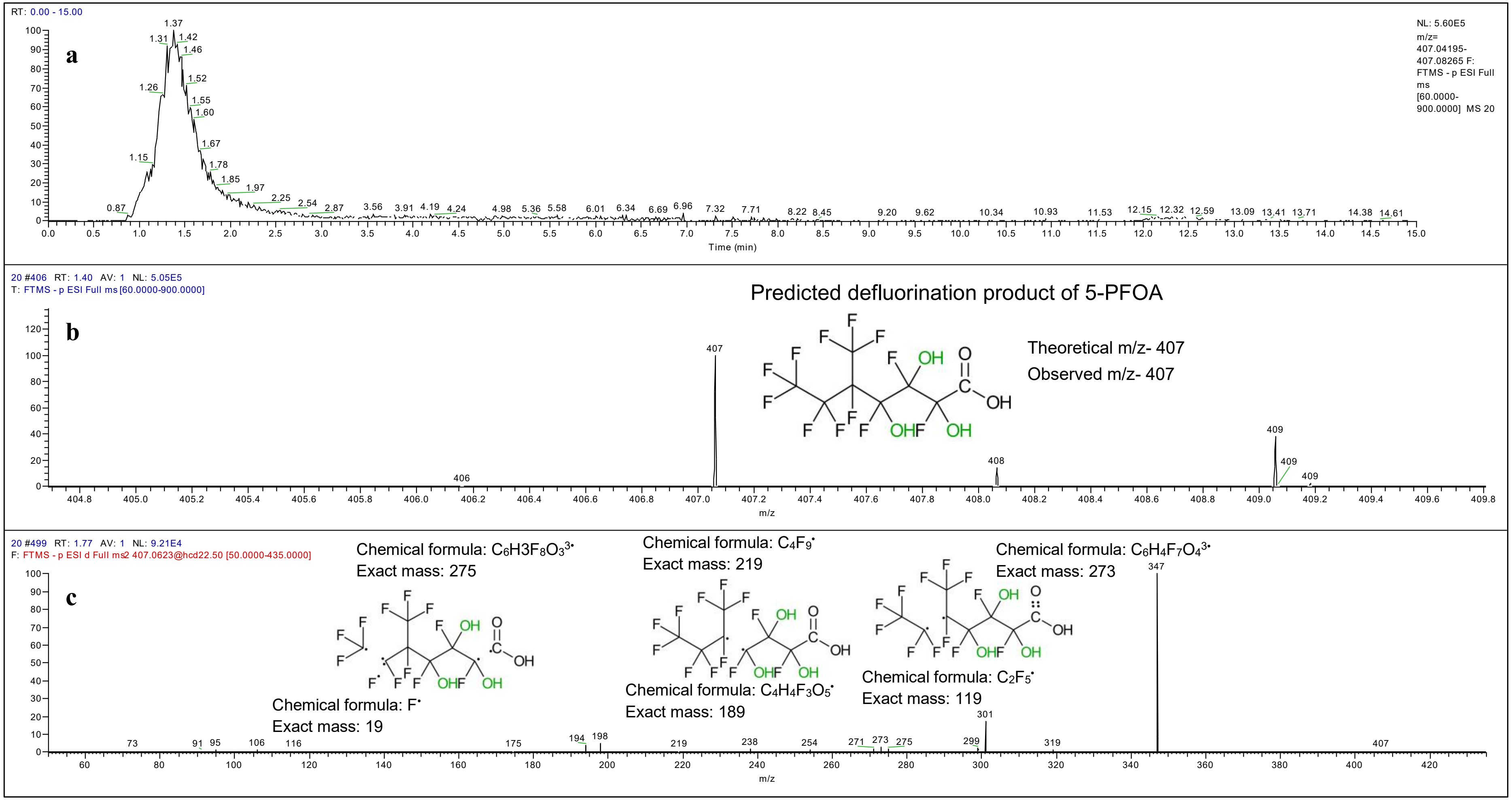

**Figure S3 LC-MS analysis of the predicted defluorination product derived from 5-PFOA.**
(**a**) Extracted ion chromatogram of the reaction product. (**b**) High-resolution mass spectrum showing the ion at *m/z* 407 corresponding to the predicted defluorination product. (c) MS/MS fragmentation spectrum of the *m/z* 407 ion, with proposed fragment assignments.

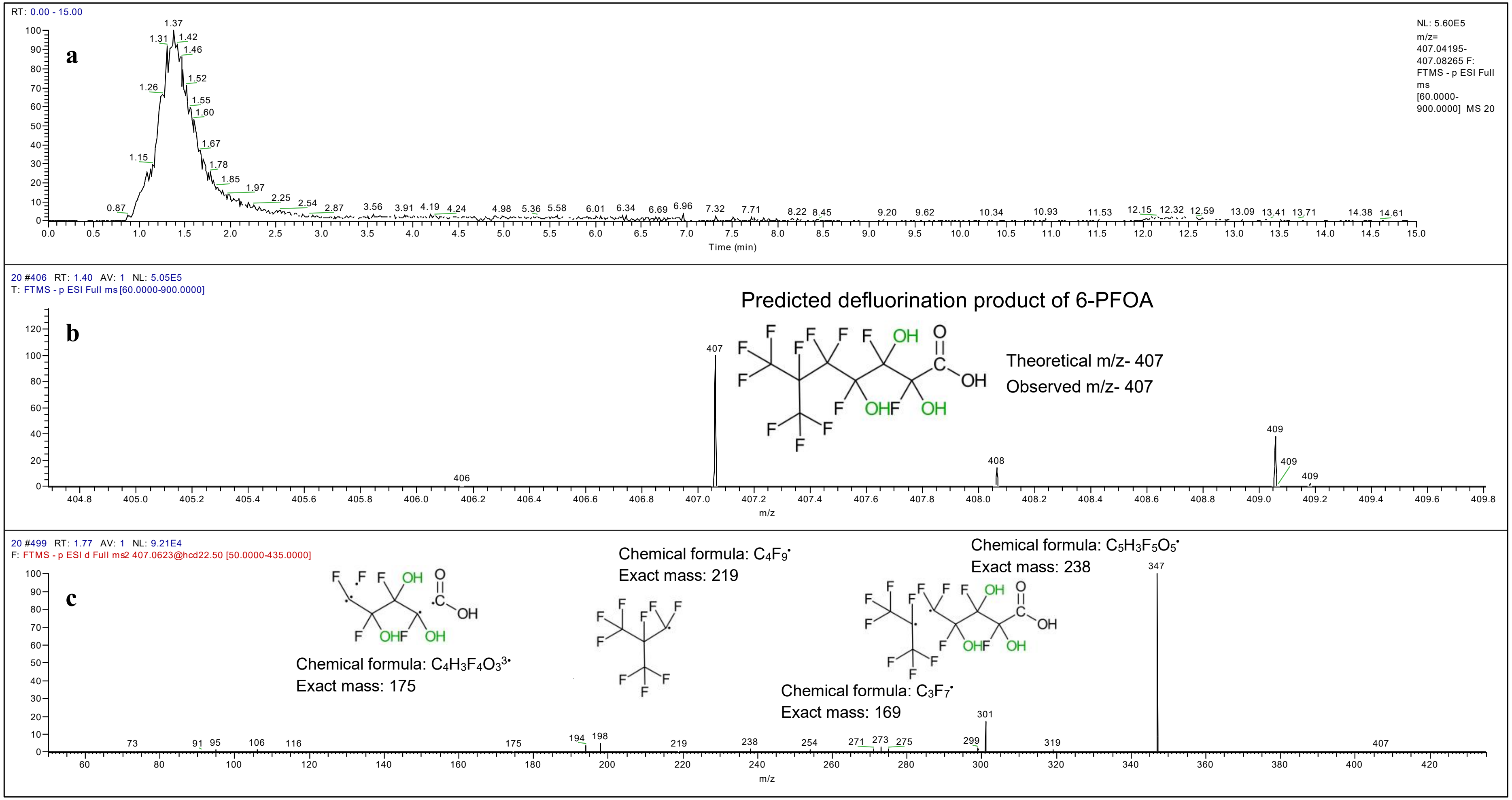

**Figure S4 LC-MS analysis of the predicted defluorination product derived from 6-PFOA.**
(**a**) Extracted ion chromatogram of the reaction product. (**b**) High-resolution mass spectrum showing the ion at *m/z* 407 corresponding to the predicted defluorination product. (c) MS/MS fragmentation spectrum of the *m/z* 407 ion, with proposed fragment assignments.

### References

(1) Harris, J. D.; Coon, C. M.; Doherty, M. E.; McHugh, E. A.; Warner, M. C.; Walters, C. L.; Orahood, O. M.; Loesch, A. E.; Hatfield, D. C.; Sitko, J. C. Engineering and characterization of dehalogenase enzymes from *Delftia acidovorans* in bioremediation of perfluorinated compounds. *Synth. Syst. Biotechnol.* **2022**, *7* (2), 671–676.

(2) Harris, J.; Gross, M.; Kemball, J.; Farajollahi, S.; Dennis, P.; Sitko, J.; Steel, J. J.; Almand, E.; Kelley-Loughnane, N.; Varaljay, V. A. Draft genome sequence of the bacterium *Delftia acidovorans* strain D4B, isolated from soil. *Microbiol. Resour. Announc.* **2021**, *10*, 10.1128/mra.00635-00621.
